# Leveraging a Comprehensive Unbiased RNAseq Database Uncovers New Human Monocyte-Derived Macrophage Subtypes Within Commonly Employed *In Vitro* Polarization Methods

**DOI:** 10.1101/2024.04.04.588150

**Authors:** Timothy Smyth, Alexis Payton, Elise Hickman, Julia E. Rager, Ilona Jaspers

## Abstract

Macrophages are pivotal innate immune cells which exhibit high phenotypic plasticity and can exist in different polarization states dependent on exposure to external stimuli. Numerous methods have been employed to simulate macrophage polarization states to test their function *in vitro*. However, limited research has explored whether these polarization methods yield comparable populations beyond key gene, cytokine, and cell surface marker expression. Here, we employ an unbiased comprehensive analysis using data organized through the **<u>a</u>**ll **<u>R</u>**NA-seq and **<u>Ch</u>**IP-**<u>s</u>**eq **<u>s</u>**ample and **<u>s</u>**ignature **<u>s</u>**earch (ARCHS^4^) database, which compiles all RNAseq data deposited into the National Center for Biotechnology Information (NCBI) Sequence Read Archive (SRA). *In silico* analyses were carried out demonstrating that commonly employed macrophage polarization methods generate distinct macrophages subsets that remained undescribed until now. Our analyses confirm existing knowledge on macrophage polarization, while revealing nuanced differences between M2a and M2c subpopulations, suggesting non-interchangeable stimuli for M2a polarization. Furthermore, we identify divergent gene expression patterns in M1 macrophages following standard polarization protocols, indicating significant subset distinctions. Consequently, equivalence cannot be assumed among polarization regimens for *in vitro* macrophage studies, particularly in simulating diverse pathogen responses.

## Introduction

Macrophages are a major innate immune cell necessary for normal development, homeostasis, and immunity^1^. Tissue resident macrophages maintain homeostasis and initiate immune responses toward pathogens^2^ while monocyte-derived macrophages (MDMs) are recruited into inflamed tissues where they serve broad pro-inflammatory and pro-resolutory functions throughout the course of an infection^3^. Macrophages can polarize to pro-inflammatory M1 or pro-resolutory M2 states depending on the particular stimuli they encounter^4,5^. Research increasingly suggests macrophages exist on a spectrum between the two states^4–6^ however, this paradigm has served as a useful model for testing macrophage responses *in vitro*. While tissue resident cells frequently stem from a unique lineage compared to MDMs, MDMs represent the major population at the site of infection through resolution of inflammation^7–9^, reinforcing the need to investigate MDMs when considering changes in macrophage function in the context of immune responses.

For mechanistic *in vitro* studies, MDMs are commonly generated from differentiated monocytes isolated from peripheral blood^10–12^. Following differentiation, resulting macrophages can be polarized to M1 states using a variety of inflammatory stimuli, but are commonly generated using lipopolysaccharide (LPS), interferon gamma (IFN-γ) or a combination of both^11–14^. Similarly, M2 macrophages can be generated using a variety of stimuli but are commonly polarized using interleukin (IL)-4,-10,-13, or a combination of IL-4 and IL-13^11–14^. While the spectrum of polarization is increasingly recognized and M2 macrophages have previously been subdivided into M2a-M2d subsets depending on polarization method and subsequent function^14^, there are not universally agreed upon methods for generation of either M1 or M2 cells, and minimal investigation into potential differences in cellular responses based on which polarization method is employed in a particular study, especially for M1-type models.

Next generation sequencing techniques, such as RNA-seq, have been employed to investigate the differences between macrophage populations, including those between polarization states. While these studies have provided a key understanding into the function of macrophages, they have generally employed limited sample sizes with singular methods for generation of each polarization state^15–17^. Despite advances in next generation sequencing techniques allowing for lower costs, large scale studies of macrophage polarization states, especially those comparing each polarization method, to determine differences in resulting macrophage populations have remained out of reach for most individual researchers. Fortunately, increased public availability of raw sequencing data allows for the higher-level integration of data collected from dozens of researchers, allowing for *in silico* investigation of these important questions.

Here, we utilized the all RNA-seq and ChIP-seq sample and signature search (ARCHS^4^) database^18^, which compiles all RNAseq data deposited into the National Center for Biotechnology Information (NCBI) Sequence Read Archive (SRA). We used the ARCHS^4^ database to investigate differences between macrophage populations, between polarization states, and within polarization states using various published polarization methods. We found that, despite longstanding acceptance of multiple methods for the generation of each polarization state, different polarization methods do not generate equivalent macrophage populations. While M2 macrophage subpopulations have been recognized by the field^14^, we found that M1 macrophages also exhibit subpopulations depending on the specific method for polarization, that these subpopulations are not equivalent, and that differences in these subpopulations should be considered when designing, analyzing, and interpreting *in vitro* macrophage studies.

## Methods

### Sample Annotation and Polarization Method Assignment

The total ARCHS^4^ version 2.2 expression (gene level) database was accessed on October 13^th^, 2023^18,19^. Sample metadata were extracted from the hdf5 data matrix into RStudio version 4.2.3^20^ using the R package rhdf5^21^ and samples were sorted based on key words. Macrophages were originally selected using ‘[M]acrophage’, ‘MDM’, and ‘[M]onocyte’ keywords, and further sorted manually to retain only primary human monocyte-derived macrophages collected from healthy control blood donors and differentiated using macrophage colony-stimulating factor (M-CSF) or granulocyte-macrophage colony-stimulating factor (GM-CSF). Samples were further sorted to remove samples treated with chemical or infectious agents based on review of the sample metadata. Samples were assigned polarization states and methods based on the treatments described in the metadata. Sample GEO accession IDs were used to extract gene level count data.

### Gene and Sample Filtering

Samples were initially selected based on polarization method to remove uncommon polarization methods such as IL-6 or treatments not of interest. Retained methods include M0, LPS, IFN-γ, LPS+IFN-γ, IL-4, IL-10, IL-13, and IL-4+IL-13. Using the R package edgeR^22^, data were organized into a DGEList object grouped by polarization method, and normalization factors were calculated. Genes were filtered for background with the edgeR function filterByExpr using default parameters to remove gene without sufficiently large counts across samples^22,23^. Normalization factors were recalculated, and values were transformed to log count per million (CPM). Sample correlations within each polarization method were calculated and samples were clustered through dendrogram clustering with a tree cut height of 0.25 (corresponding to a Pearson correlation of ≥ 0.75). To detect and remove potential sample outliers, the largest dendrogram cluster for each polarization method was selected and the remaining samples not sorted into this cluster were removed. Untransformed sample selected count data were saved for differential expression analysis while log CPM data were corrected for batch effect using the limma^24^ function removeBatchEffect with GEO series ID acting as batch designation. Batch correction serves to correct for variation in the data caused by non-biological effects such as individual technician handling, reagents, and variation in instrumentation^25^. Batch corrected data were saved for statistical testing and downstream analysis.

### Differential Expression using Limma-Voom

Differential expression between samples was conducted using limma-voom^26^ in accordance with previously described methods^27,28^. Briefly, raw count data were assembled into a DGElist object and grouped by polarization method. Normalization factors were calculated to scale the raw library sizes into effective library sizes, and a model matrix was created accounting for polarization state or polarization method and GEO series ID to account for batch effects. The limma function voom was performed to calculate the mean-variance relationship of the log-counts and generate a precision weight for each observation^26^ before fitting gene-wise linear models for each treatment factor. Contrasts of interest were estimated for each gene, empirical Bayes smoothing was applied to gene-wise standard errors^29^, and resulting statistics were presented with the limma function topTable. Samples were considered significantly differentially expressed if Benjamini-Hochberg (BH)-adjusted P-values were less than 0.05 and log2 fold changes were greater than or equal to 2, or less than or equal to -2. Volcano plots were generated using the resulting data using the EnhancedVolcano R package^30^. Euler plots of overlapping DEGs between comparisons were assembled using the R package eulerr^31^. Differentially expressed genes from comparisons of interest were analyzed using Qiagen Ingenuity Pathway Analysis (IPA, QIAGEN Inc.) and the resulting data were imported into RStudio. The top canonical pathway results as determined by IPA were extracted and ranked by p-value and the top 10 canonical pathways of each comparison were extracted. Z-scores of the resulting pathways, which describes positive or negative changes in pathway activity from each comparison, were compiled and presented as a heatmap using the R package ComplexHeatmap^32^.

### Random Forest Modeling

The supervised machine learning algorithm random forest^33^ was employed to determine whether macrophages could be sorted between polarization methods and to determine the key parameters linked to each polarization method. Random forest modeling is an ensemble machine learning method which constructs multiple decision trees based on a subset of input features, and for classification analysis, classifies samples based on majority voting of the resulting decision trees^33,34^. Batch corrected samples were subsetted to retain M0, IFN-γ, LPS, and IFN-γ+LPS polarization methods and assessed for statistical differences between polarization methods using One-Way ANOVA. Genes with BH (FDR) adjusted p-values below 0.05 were retained and used for modeling. Samples were split 70/30 into a training and testing data subset. Bootstrap aggregation (bagging) was employed to randomly select samples for fitting to decision trees^33,34^. To balance the likelihood of selecting underrepresented polarization methods, the number of samples in each group (polarization method) of the training data set was measured and sample case weights were calculated based on these numbers. Briefly, the fraction of each group to the whole sample number was calculated and a sampling ratio of the whole was calculated. The random forest was grown using the R package ranger^34^ with 1500 trees, a max tree depth of 15 splits, the number of features (genes) assessed at each split (mtry) equal to the square root of the gene number, and case weights set to the sampling ratio of the whole. Model out of bag (OOB) error rate, which measures the percentage of misclassified samples from the training data not included in the bootstrapped data set, was assessed, and model performance was assessed using confusion matrices which describe the model’s ability to correctly classify samples to their known polarization method. In addition, model permutation importance was assessed to determine the impact of each feature on the performance of the model by comparing OOB error rates for an unpermuted feature to OOB error rates after permuting that feature across the OOB data set. The mean difference is calculated for the whole forest to achieve the permutation importance value of each feature^33,34^. Genes were ranked based on descending permutation importance and the top 10,000 genes were retained. The training and testing data set were reduced to these 10,000 genes and the model was retrained with the same specifications as above. Model OOB rate, permutation importance, and performance indices were assessed. After recalculating permutation importance of the model, the top 1,000 genes by permutation importance were isolated and the training and testing data set were again reduced to these 1,000 genes.

### Sample PCA Clustering and Heatmap Generation

The top 1,000 genes by permutation importance as determined by random forest modeling were used for sample principal component analysis (PCA) clustering and heatmap generation. PCA is a multivariate statistical technique which reduces the dimensionality of data by identifying a set of new orthogonal variables called principal components along which the variation in the data is maximized. This allows for the representation of thousands of variables by a relatively low number of new variables which can be easily plotted and assessed for similarities and differences^35,36^. Briefly, batch corrected log CPM count data for the 1,000 selected genes were isolated and PCA was performed using the R package factoextra^37^. Partition around medoid (PAM) clustering^38^ (K = 4 clusters) of genes and samples by polarization method were calculated from scaled data using the R package parallelpam^39^. Heatmaps were generated using scaled log CPM count data using the R package ComplexHeatmap clustering rows (genes) and columns (polarization method) using the PAM clustering assignments.

### Gene Ontology and Gene Set Variation Analysis

Gene ontology (GO) terms were assigned to genes retained following filterByExpr (gene universe) using the R package biomaRt^40^. Genes present in the top 1000 genes by random forest permutation importance were tested for GO biological process (BP) term enrichment against the gene universe using the R package topGO^41^ with the weight01 algorithm and fisher statistic. GO term enrichment tests for significantly increased presence of genes linked to a particular GO term within a gene list compared to the background gene universe^42^. Genes contained within significantly enriched GO BP terms were assembled as gene sets using the R package GSEABase^43^ and gene set variation analysis was performed using the R package GSVA^44^. GSVA is a gene set enrichment analysis method which calculates sample-wise enrichment scores for a whole gene set as a function of genes inside and outside the gene set and estimates variation of gene set enrichment over samples independently of group designations^44^. Differential expression of resulting GSVA enrichment scores was performed using the R package limma with M0 samples serving as the reference group. Resulting comparisons were arranged by ascending BH adjusted p-value and the top 20 pathways of each polarization method were selected. Differential expression of GSVA enrichment scores were calculated between groups of interest and BH adjusted p-values were retained. GSVA enrichment scores or pathway statistical comparisons were presented as heatmaps using the R package ComplexHeatmap.

### Random Forest Prediction of Pathogen Exposed Samples

Monocyte-derived macrophage samples initially removed due to exposure to infectious agents were resorted and assigned infectious agent information, retaining samples from groups with at least 5 samples per group. Genes were removed, samples were selected, and batch effects were corrected in accordance with the original dataset. The previously trained random forest models were used to predict M0, LPS, IFN-γ, or LPS+IFN-γ states for infectious agent exposed samples. Results of the prediction for each model were graphed using the R package ggplot2^45^.

## Results

### Monocyte-derived Macrophages Demonstrate Distinct Gene Expression Patterns between Polarization States and Polarization Methods

ARCHS4 version 2.2 was downloaded and extracted, resulting in a dataset with 722,425 samples with 67,186 genes. Data associated with 1,444 primary human monocyte-derived macrophage (MDMs) samples were extracted and assigned polarization information based on a combination of metadata labels and manual annotations. Following sample filtering using dendrogram clustering (see Methods), a total of 1,075 samples were retained (Table 1). DE analysis was performed using limma-voom^26^ between either sample polarization state (Figure 1) or polarization method (Figure 2, 3) with Benjamini-Hochberg (BH) adjusted p-values < 0.05 and absolute log2 fold changes ≥ 2 being considered significantly differentially expressed.

**Table 1.** Gene and sample numbers before and after filtering. Genes were filtered using the edgeR function filterByExpr. Samples were filtered using dendrogram clustering with cut heights of 0.25 corresponding to a Pearson correlation ≥ 0.75.

|  | Unfiltered | Filtered |
| --- | --- | --- |
| M0 | 949 | 713 |
| M1 | 367 | 260 |
| M2 | 128 | 102 |
| IFN | 48 | 47 |
| LPS | 106 | 102 |
| IFN_LPS | 213 | 111 |
| IL4 | 51 | 51 |
| IL10 | 25 | 20 |
| IL13 | 7 | 7 |
| IL4_IL13 | 45 | 24 |
| Genes | 67186 | 40545 |

**Figure 1.**
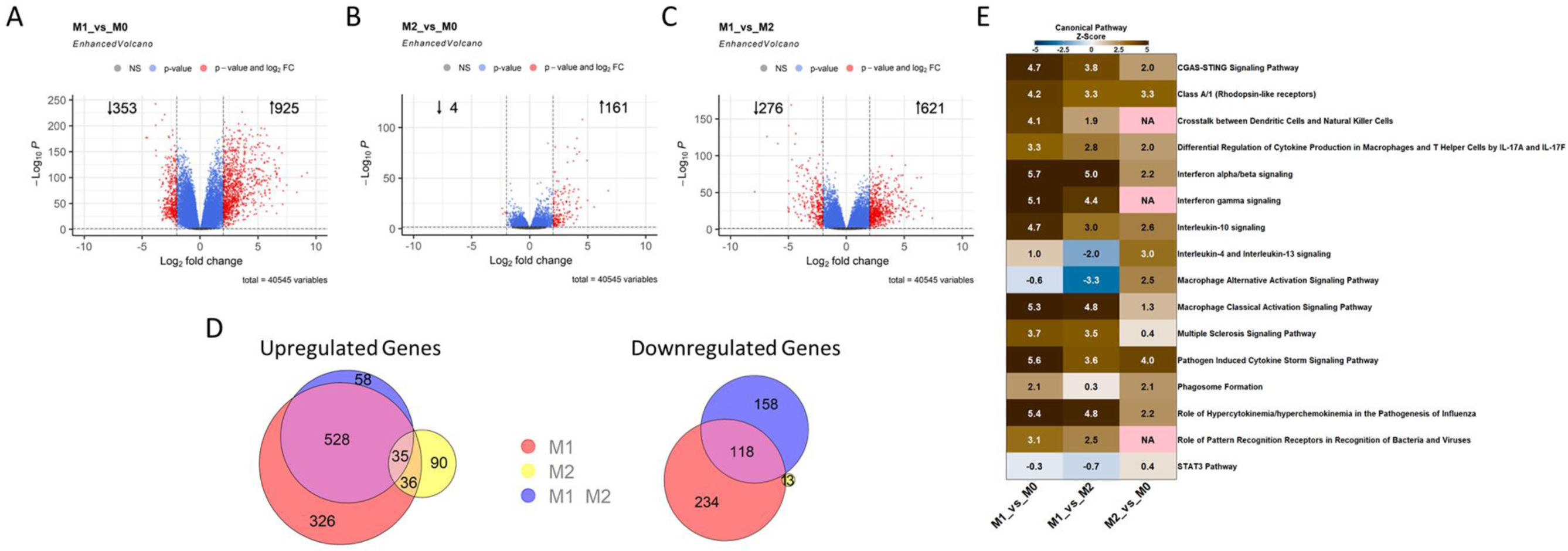
Human Monocyte-Derived Macrophages Demonstrate Clear Polarization-Specific Gene Expression Patterns. A-C. Volcano plots representing results of differential expression analyses compared to unpolarized M0 macrophages (A-B) or between M1 and M2 macrophages (C). D. Euler plots describing overlap of significantly differentially expressed genes between groups. E. Ingenuity Pathway Analysis (IPA) canonical pathway activation z-score heatmap. Significance cutoffs of Benjamini-Hochberg (BH)-adjusted P-values < 0.05 and absolute log2 fold changes ≥ 2 were employed for all analyses.

**Figure 2.**
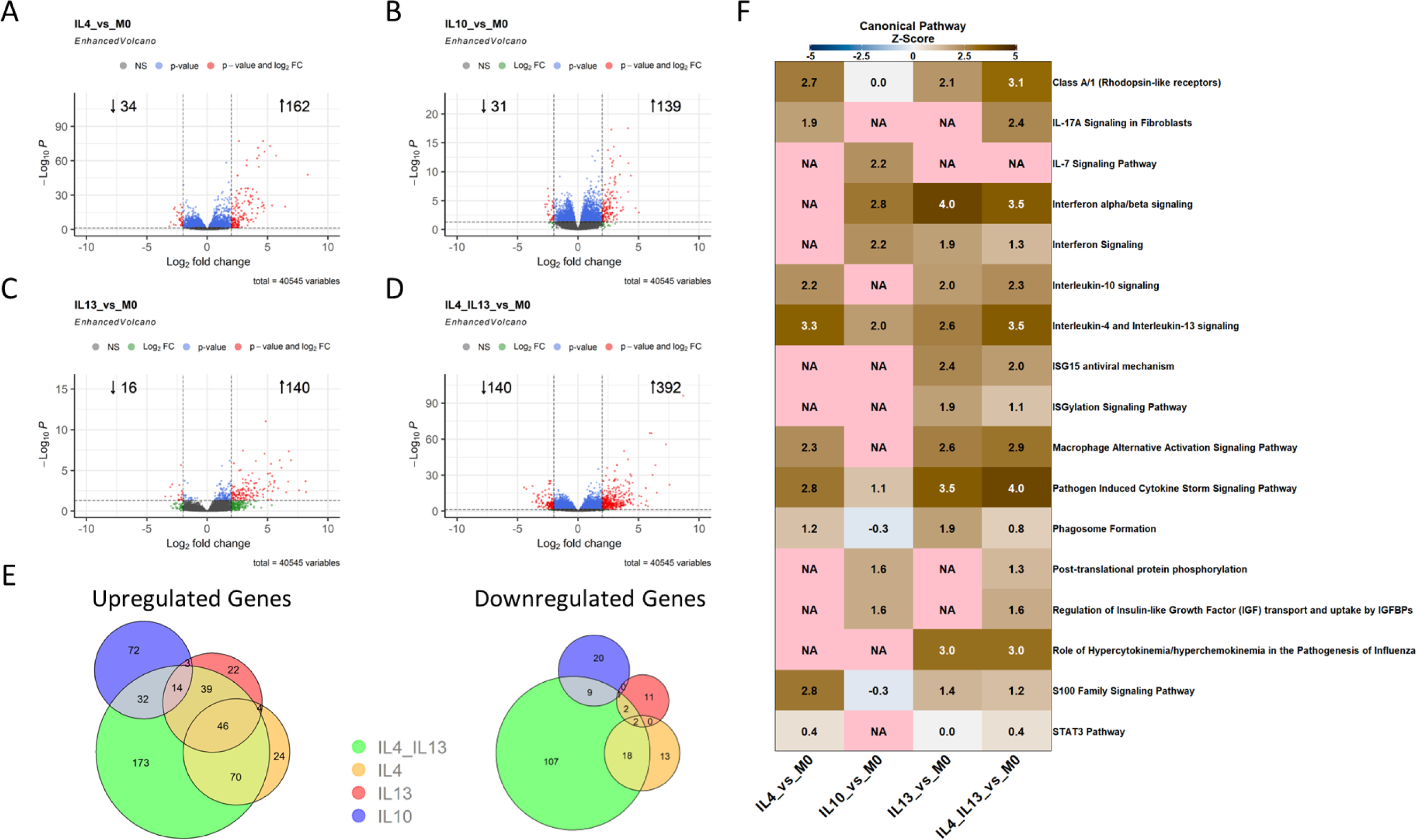
M2 Macrophages Demonstrate Polarization Method-Dependent Gene Expression Patterns. A-D. Volcano plots representing results of differential expression analyses compared to unpolarized M0 macrophages. E. Euler plots describing overlap of significantly differentially expressed genes between groups. F. Ingenuity Pathway Analysis (IPA) canonical pathway activation z-score heatmap. Significance cutoffs of Benjamini-Hochberg (BH)-adjusted P-values < 0.05 and absolute log2 fold changes ≥ 2 were employed for all analyses. NA values represent pathway z-scores which could not be calculated.

**Figure 3.**
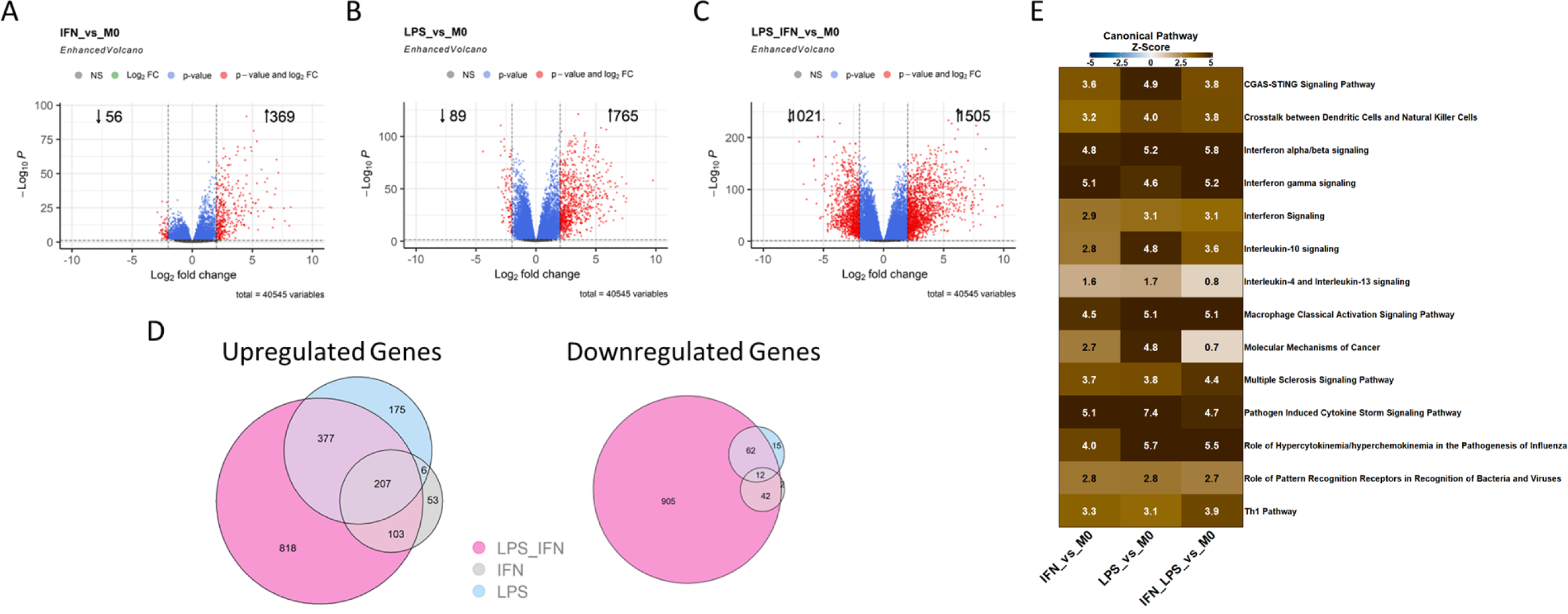
M1 Macrophages Demonstrate Distinct Gene Expression Patterns Between Polarization Methods. A-C. Volcano plots representing results of differential expression analyses compared to unpolarized M0 macrophages. D. Euler plots describing overlap of significantly differentially expressed genes between groups. E. Ingenuity Pathway Analysis (IPA) canonical pathway activation z-score heatmap. Significance cutoffs of Benjamini-Hochberg (BH)-adjusted P-values < 0.05 and absolute log2 fold changes ≥ 2 were employed for all analyses.

DE analysis between polarization state demonstrated clear differences between M1 populations compared to M0 and M2 populations with 1,278 (Figure 1A) and 897 (Figure 1C) DEGs, respectively, and a lower degree of differences detected between M2 and M0 populations with 165 DEGs (Figure 1B). Significant overlap was detected between comparisons with 646 genes found to be differentially expressed in M1 macrophages compared to both M0 and M2 macrophages; however, 560 genes are uniquely differentially expressed when M0 macrophages serve as the reference group, while 216 genes are uniquely differentially expressed when M2 macrophages serve as the reference group (Figure 1D). Following Ingenuity Pathway Analysis (IPA), clear pathway patterns were detected, with M1 cells inducing pro-inflammatory and classically activated macrophage pathways while M2 cells demonstrated reduced activity of these pathways, increased IL-4/IL-13 signaling, and alternatively activated macrophage pathways (Figure 1F). Together, these findings suggest this data set agrees with previously demonstrated results from smaller scale studies and can serve as a useful model for examining more specific polarization method-specific findings.

### IL-4/IL-13 Polarized M2 Macrophages Demonstrate Distinct Gene Expression Patterns Compared to IL-10 Polarized M2 Cells

Initial characterization of M2 macrophages segregated by polarization method suggests gene expression patterns compared to unpolarized M0 macrophages are largely similar (Figure 2). Co-exposure to IL-4+IL-13 demonstrated the greatest change in gene expression patterns with both the highest number of DEGs as well as the greatest number of unique DEGs (Figure 2D/E). Single exposures to IL-4 or IL-13 induced similar gene expression patterns to co-exposures with overall lower DEG numbers (Figure 2A, C-E); however, unique DEGs were detected in each group (Figure 2E). IL-10 exposure induced a larger number of unique DEGs to single or co-exposure to IL-4 and/or IL-13 (Figure 2E). When considering canonical pathway activity, a large degree of overlap was detected between IL-4, IL-13, and IL-4+IL-13 patterns, while IL-10 exposed macrophages demonstrated minimal overlap with IL-4 and slight overlap with either IL-13 or IL-4+IL-13 exposed macrophages (Figure 2F). For example, the genes associated with the IL-7 signaling pathway were only differentially expressed in IL-10 exposed macrophages (Figure 2F). We also sought to directly assess differential expression between polarization methods. Slight differences in gene expression patterns were detected with the results of these comparisons presented in Supplemental Figure S2.

### M1 Polarization Methods Induce Previously Undescribed Macrophage Subpopulation

Distinct gene expression profiles were detected between IFN-γ, LPS, and LPS+IFN-γ co-exposed macrophages to an even greater extent than M2 macrophage subsets (Figure 3). As with M2 groups, initial characterization of M1 macrophages segregated by polarization method demonstrated clear differences in gene expression patterns compared to unpolarized M0 macrophages in the number of total and unique differentially expressed genes and the magnitude of canonical pathway activation. Specifically, IFN-γ exposed macrophages demonstrate the lowest number of unique and total DEGs (Figure 3A, Figure 3D). LPS exposure induced greater numbers of total and unique DEGs compared to IFN-γ exposure (Figure 3B, 3D), while co-exposure to LPS+IFN-γ induced the greatest number of total and unique DEGs of all groups (Figure 3C, 3D). While overlap in DEGs is detected between all three groups, more DEGs are distinct to a single polarization method than are shared between polarization methods (Figure 3D). Canonical pathway activation followed similar directionality between each polarization condition; however, clear differences are detected in the magnitude of activation between polarization methods (Figure 3E). For example, genes associated with Molecular Mechanisms of Cancer pathway show clear differences between the three M1 macrophage groups (Figure 3E). As with M2 macrophages, we sought to directly compare M1 polarization methods to each other to determine differential expression between groups. Major differences were detected between polarization methods, and results of these comparisons are presented in Supplemental Figure S3.

### Random Forest Modeling Describes Key Genes Corresponding to Each Macrophage Population following Exposure to Traditional M1-Polarizing Stimuli

As IFN-γ, LPS, and LPS+IFN-γ polarized macrophages demonstrated clear differences in gene expression profiles and canonical pathway activity, we sought to determine whether macrophages could be accurately classified by polarization method using random forest modeling^33,34^ (see Methods), and to determine the genes most impactful in producing classification predictions. Following initial modeling (Supplemental Figure S4A), genes were ranked based on descending permutation importance (see Methods). Following ranking, the model was retrained using the top 10,000 (Supplemental Figure S4B) and then top 1,000 genes by permutation importance, resulting in a model with an overall accuracy of 0.98 (Figure 4). The model demonstrated overall high sensitivity, specificity, and precision, with IFN-γ exposed cells demonstrating the lowest sensitivity at 0.89 and LPS demonstrating the lowest precision at 0.87 (Figure 4). Out of bag (OOB) error rates for the model improved from 1.03% to 0.88%, ending at 0.59% when the model was limited to the top 1,000 genes (data not shown).

**Figure 4.**
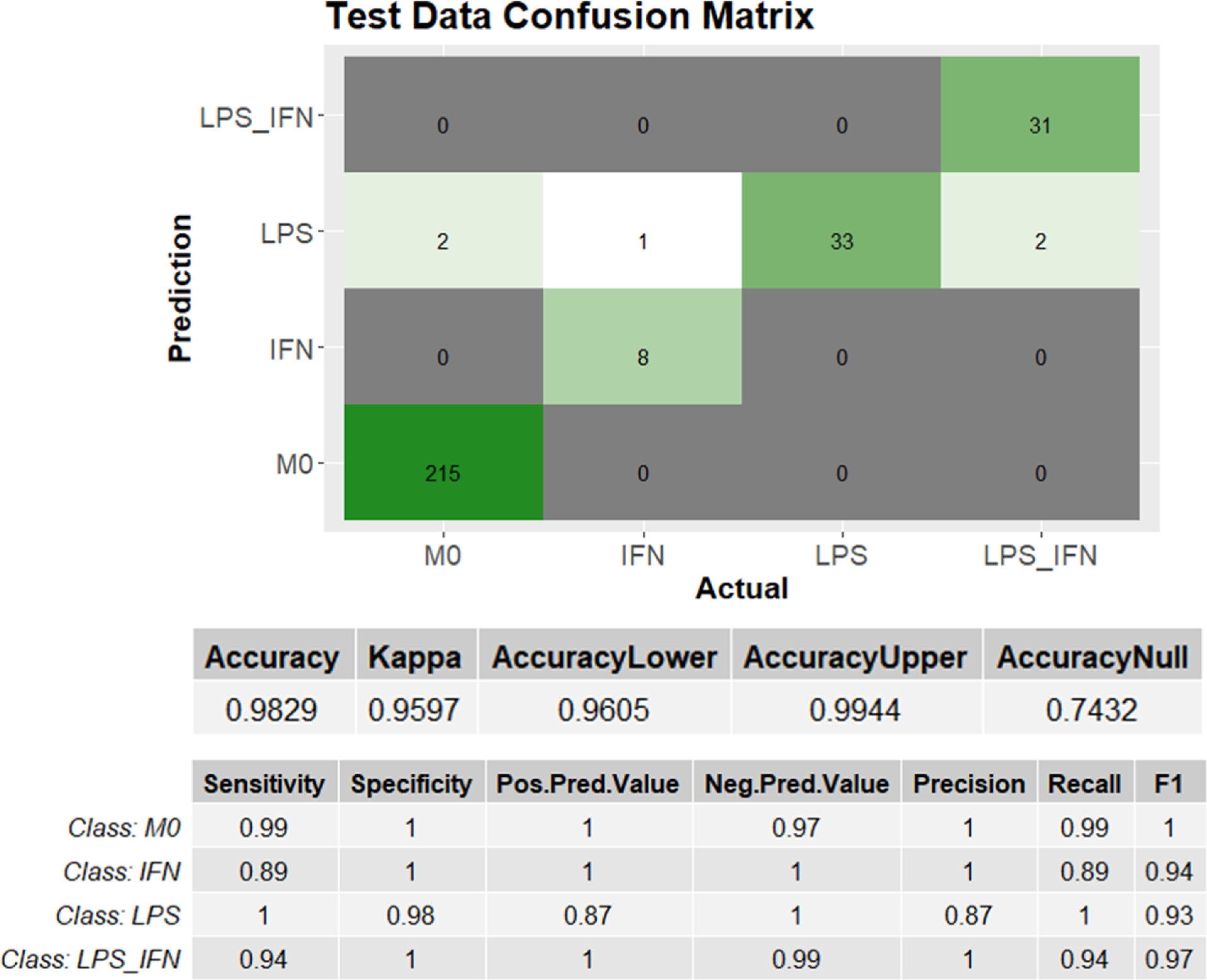
Random Forest Demonstrates High Model Performance in Classification of M1 Macrophage Subpopulations. Random forest confusion matrix (top) describing the number of samples from the test data subset predicted as each class versus actual class. Model performance metrics (middle, bottom) describe the ability of the model to correctly classify samples and to avoid misclassifying samples across classes.

Visualization of the top 1,000 genes further demonstrates differences between M1 macrophage subtypes. Partition around medoid (PAM) clustering reveals macrophage subtypes sort well according to the defined polarization stimuli (Figure 5A). Specifically, 99.4% of samples sorted into cluster 1 were M0 cells, 95.4% of samples in cluster 2 were LPS+IFN-γ exposed, 92.2% of samples in cluster 3 were LPS exposed, and 86.8% of samples in cluster 4 were IFN-γ exposed (Figure 5A). LPS+IFN-γ exposed cells demonstrate the overall highest gene expression as measured by row-scaled z-scores (Figure 5A). Principle component analysis (PCA) further demonstrates differences between gene expression patterns of M1 subtypes (Figure 5B). More overlap is detected between cells singly exposed to IFN-γ or LPS as demonstrated by both PAM clustering (Figure 5A) and PCA clustering (Figure 5B); however, each method demonstrates distinct patterns, supporting the conclusion that polarization method induces a distinct macrophage subpopulation.

**Figure 5.**
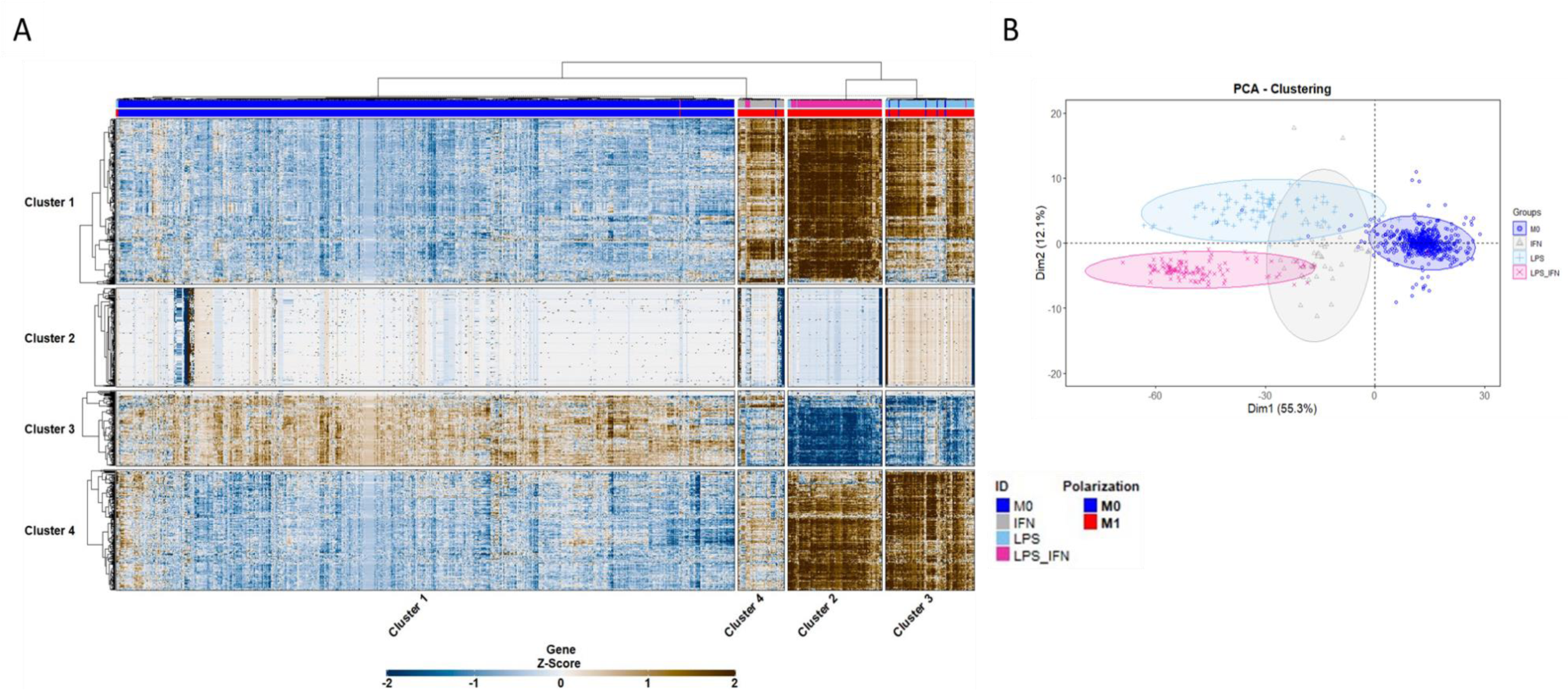
M1 Macrophage Subpopulations Demonstrate Distinct Gene Expression Patterns. A. Heatmap representing row-scaled z-scores of log count per million (CPM) reads for the top 1000 genes as determined by random forest permutation importance. Rows (genes) and columns (samples) were clustered using partition around medoid (PAM) clustering with k=4 medoids. B. Principal component analysis (PCA) plot with concentration ellipses demonstrating clustering by defined polarization method.

### Gene Set Variation Analysis Demonstrates Polarization Method-Dependent Patterns

To further investigate the differences in gene expression patterns demonstrated through PAM and PCA clustering, gene set variation analysis (GSVA), a gene set enrichment analysis method which calculates sample-wise enrichment scores for a whole gene set^44^ (see Methods), was performed. Gene sets were selected and assembled based on significantly enriched gene ontology biological process (BP) terms within the top 1,000 genes described in Figure 5, and GSVA was performed on the 40,545 selected genes comparing all M0, LPS, IFN-γ, and LPS+IFN-γ samples. Consistent with our previous findings, distinct differences were detected between each macrophage subpopulation. LPS and LPS+IFN-γ exposed macrophages demonstrated more similar pathway activity compared to IFN-γ exposed macrophages (Figure 6A); however, most pathways were still found to be significantly differentially expressed between each polarization method, particularly those involving cellular responses to cytokines and chemokines (Figure 6B). Overall, pathway differential expression in LPS+IFN-γ exposed macrophages was consistent when either LPS or IFN-γ served as the reference group (Figure 6B). However, several key pathways involved in the response to non-viral agents were similar between co-exposure and LPS exposure alone, while pathways involved in the response to viral agents were similar in co-exposed cells compared to cells exposure to IFN-γ alone (Figure 6B), suggesting co-exposed macrophages retain these key characteristics commonly attributed to LPS or IFN-γ. Overall, these findings further support the notion that each polarization method induces a unique population of macrophages.

**Figure 6.**
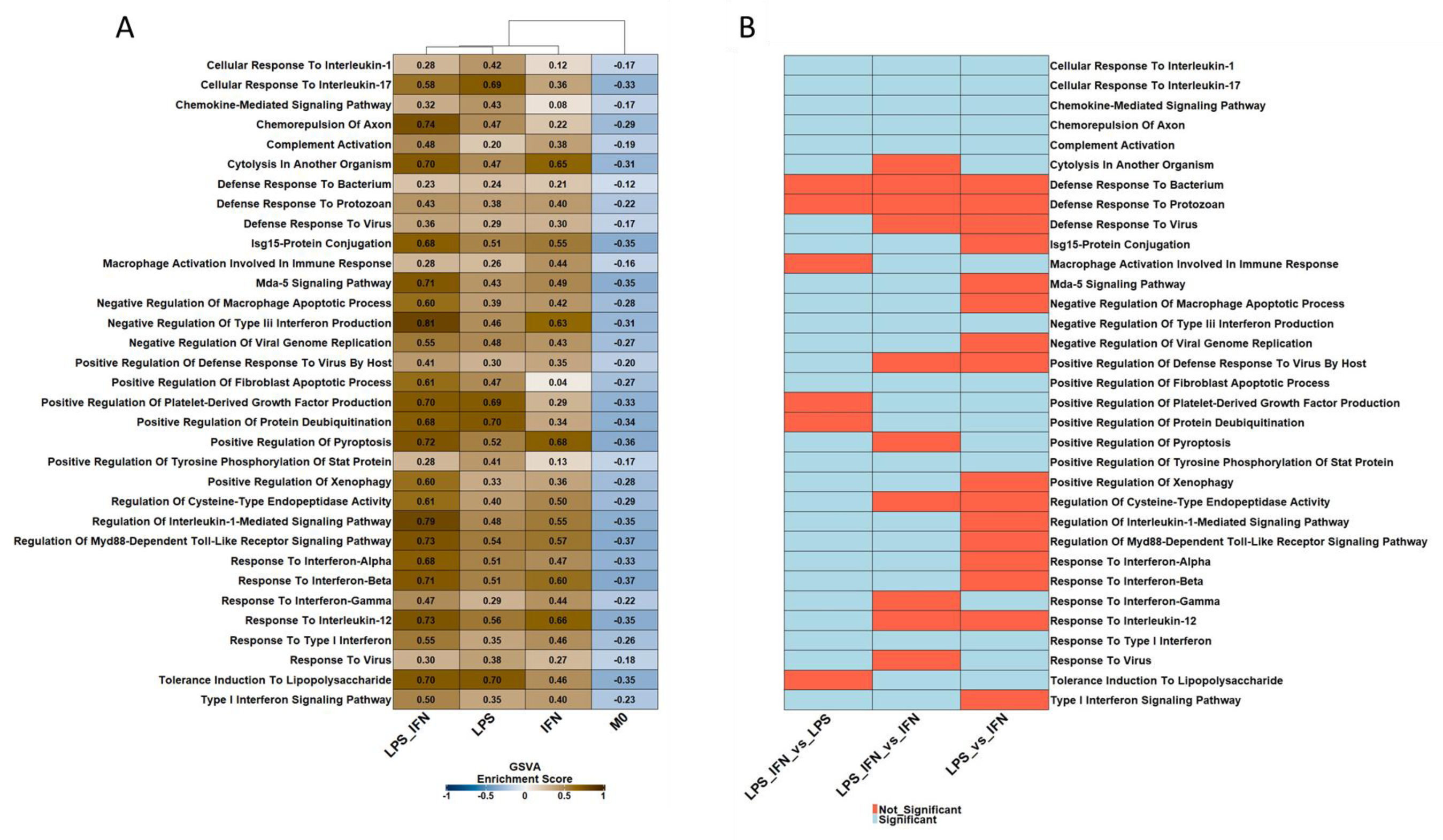
M1 Macrophage Subpopulations Demonstrate Varied Gene Ontology Biological Pathway Activity. A. Heatmap of mean gene set variation analysis (GSVA) enrichment scores by indicated polarization method. Presented pathways represent the top 20 most significantly differentially expressed pathways compared to M0 macrophages following GSVA analysis of all significant enriched gene ontology biological process terms from the top 1000 genes as determined by random forest permutation importance. B. Heatmap representing results of differential expression analysis of GSVA enrichment scores between indicated groups.

### IFN-γ Polarization Induces a Poor Model for Bacterially or Virally Infected Macrophages

Data associated with monocyte-derived macrophage (MDM) samples in the ARCHS^4^ database exposed to bacterial or viral agents were extracted and processed in accordance with previously described methods. Infectious agents and sample numbers are presented in Table 2. Samples were classified as M0, IFN-γ, LPS, or LPS+IFN-γ polarized macrophages using predictions from the random forest models described in Supplemental Figure S4 and Figure 4. Surprisingly, regardless of the version used, random forest modeling suggests gene expression in infected MDM samples did not resemble IFN-γ polarized macrophages, but instead resemble either unpolarized M0 cells or LPS+IFN-γ exposed macrophages (Figure 7). Moreover, when random forest models are built using subsequently fewer genes which were determined to hold greater permutation importance, *Listeria* and *M. tuberculosis* exposed samples increasingly resemble macrophages exposed to LPS alone, suggesting expression of the most impactful genes in these infection paradigms are likely LPS driven (Figure 7B, 7C). However, *L. pneumophila, M. smegmatis, S. epidermidis, Y. pseudotuberculosis,* and the largest sample group, *S. typhimurium,* largely retained their original prediction of LPS+IFN-γ, suggesting different pathogens are capable of inducing gene expression patterns similar to either single exposures to LPS or co-exposure to both LPS+IFN-γ. Interestingly, HIV exposed macrophages, and to a lesser extent, Zika virus exposed macrophages, resemble LPS+IFN-γ exposed cells, compared to other virally infected cells which mostly resemble M0 macrophages regardless of the number of genes retained in the model (Figure 7). Together, these findings suggest IFN-γ is not an appropriate model to reflect infected macrophages, that LPS exposure alone only reflects macrophages of a subset of bacterial exposures, and that co-exposure to LPS+IFN-γ is likely the most applicable model for replicating macrophages exposed to bacterial agents.

**Table 2.** Numbers of pathogen exposed monocyte-derived macrophage samples before and after filtering.

| Agent | Unfiltered | Filtered |
| --- | --- | --- |
| Salmonella typhimurium | 268 | 268 |
| Listeria | 232 | 232 |
| Mycobacterium tuberculosis | 154 | 144 |
| HIV | 40 | 40 |
| IAV | 33 | 33 |
| Zika Virus | 33 | 33 |
| Yersinia pseudotuberculosis | 18 | 18 |
| Mycobacterium smegmatis | 17 | 17 |
| Staphylococcus epidermidis | 13 | 12 |
| Ebola Virus | 9 | 9 |
| Reston Virus | 9 | 9 |
| Chikungunya Virus | 8 | 8 |
| L. pneumophila | 6 | 6 |

**Figure 7.**
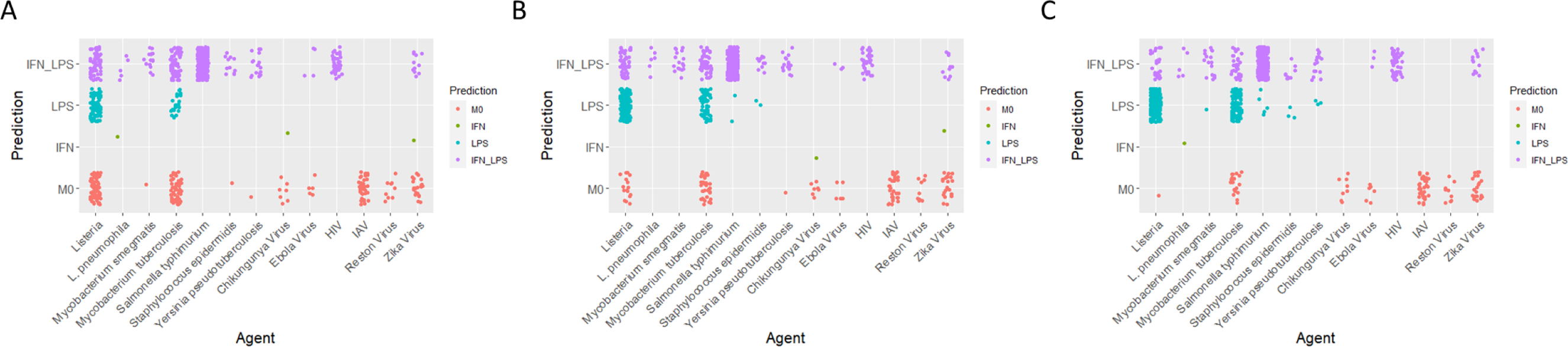
IFN-γ Represents a Poor Method for Modeling Pathogen Exposed Monocyte-Derived Macrophages. Random forest predictions of macrophage polarization methods for human monocyte-derived macrophages exposed to indicated pathogens. A. Predictions from the total model. B. Predictions from the model generated using the top 10000 genes by permutation importance of the total model. C. Predictions from the model generated using the top 1000 genes by permutation importance of the model described in panel B.

## Discussion

Here, we leveraged the ARCHS^4^ database to conduct, to our knowledge, the largest RNAseq analysis of monocyte-derived macrophage polarization states to date. This dataset reflects conclusions previously made regarding general polarization state gene expression patterns and further characterizes the IL-4/IL-13 polarized M2a and IL-10 polarized M2c subsets. More importantly, we identified previously undescribed M1 subsets corresponding to the use of LPS, IFN-γ, or LPS+IFN-γ to induce differential polarization with unique phenotypes. These findings suggest that, like M2 macrophages, M1 macrophages exist as several distinct subpopulations, demonstrating drastically different gene expression profiles, and that accepted M1 polarization and phenotyping methods cannot be considered equivalent for *in vitro* studies.

Macrophages are phagocytic innate immune cells first described by Metchnikoff in the late 1800s^46^. Metchnikoff described macrophage activation in 1905 following observations of increased bacterial killing properties of macrophages from infected animals^47^, and this observation of macrophage activation expanded to the concept of macrophage polarization, whereas macrophages are modified following exposure to various extracellular stimuli leading to a diversity of function^4,5^. Macrophage polarization was quickly sorted into the pro-inflammatory M1 and pro-resolutory M2 state in a similar manner to that applied to Th1 and Th2 lymphocytes^48^. Using this classification system, we grouped our samples into unpolarized M0, pro-inflammatory M1, and pro-resolutory M2 polarization groups and investigated differential gene expression in order to compare our model to previously published findings. In agreement with these previous findings^49–52^, M1 and M2 macrophages demonstrated large scale differential expression compared to M0 macrophages and to each other (Figure 1A-D). Further, when M0 macrophages served as the reference group, M1 macrophages demonstrated a greater number of differentially expressed genes (DEGs) compared to M2 macrophages (Figure 1A-D)^49,52^. M1 and M2 macrophages also demonstrated clear differences in gene expression patterns as shown by nearly 900 DEGs between each other (Figure 1C), which has again been suggested previously^49,50^. When considering canonical pathway modifications within each polarization group, our findings were in line with those previously reported by other researchers (Figure 1E). Namely, M1 macrophages demonstrated greater classical activation signaling pathways while M2 macrophages demonstrated greater alternative activation signaling pathways (Figure 1E). In addition, M1 macrophages demonstrated a clear upregulation of the Multiple Sclerosis (MS), interferon alpha/beta, and interferon gamma signaling pathways compared to both M0 and M2 macrophages (Figure 1E). As M1 macrophages have been linked to MS progression^53–55^ and elevated interferon-induced gene signatures have been demonstrated in peripheral blood cells of MS patients^56^, these findings further support the use of these macrophage models in investigating clinically relevant diseases as well as the use of this dataset for investigation of macrophage polarization states. This suggests the ARCHS^4^ dataset supports previously described macrophage polarization and can serve as a data source to further investigate differences in gene expression patterns within macrophage polarization states.

While the binary M1-M2 classification system has largely persisted through time, research increasingly demonstrates that macrophages exist on a spectrum between these two extremes based on a mixture of cellular exposures, signals from the immediate cellular milieu, and biological activities/function^4–6^. With this understanding, M2 macrophages have been identified to exist within subpopulations exhibiting distinct functions, termed M2a-M2d^57–59^. M2a macrophages are most in line with the traditional ideal of a pro-resolutory macrophage, increasing secretion of anti-inflammatory and immune regulatory cytokines, endocytosis, as well as tissue repair and are generated through the exposure to IL-4 and/or IL-13^57–59^. In contrast, M2c macrophages demonstrate general inactivity and play a major role in phagocytosis of apoptotic cells being generated through exposure to IL-10, glucocorticoids, and TGF-β^58,59^. Our findings related to the use of IL-4 and/or IL-13 versus IL-10 for the generation of M2 macrophages are largely in line with previously published results. IL-10 polarized macrophages demonstrated a distinct gene expression pattern compared to IL-4, IL-13, or IL-4+IL-13 polarized cells as demonstrated by gene expression patterns and canonical pathway activity (Supplemental Figure S2, Figure 2). Interestingly, while M2c macrophages have been associated with increased phagocytic activity^58,59^, IL-10 treated macrophages demonstrated negligible increases in genes associated with phagosome formation compared to M0 macrophages (Figure 2F) while demonstrating increased gene expression changes related to the IL-7 Signaling pathway activity compared directly to IL-4, IL-13, or IL-4+IL-13 (Figure 2F; Supplemental Figure S2H). However, in agreement with previous results^58,59^, IL-10 exposed macrophages demonstrate overall lower activity with generally lower canonical pathway activation (Figure 2F, Supplemental Figure S2H). In contrast to previous thought, IL-4, IL-13, or IL-4+IL-13 are not entirely equivalent polarization methods for the generation of M2a macrophages. While IL-4, IL-13, and the combination of IL-4+IL-13 each induce differential expression compared to M0 cells, each group induces DEGs not induced by other polarization methods, with the greatest number of unique DEGs found in IL-4+IL-13 co-exposed cells (Figure 2). When polarization methods are compared to each other, IL-10 continues to demonstrate a clear difference to each traditional M2a stimuli as demonstrated by unique DEGs (Supplemental Figure S2G) and canonical pathway activity including reduced macrophage alternative activation signaling pathways, IL-13 signaling pathways, and elevated phagosome formation (Supplemental Figure S2H). When directly compared, IL-4+IL-13 polarized cells demonstrate greater differences to IL-4 than IL-13 polarized cells as measured by unique DEGs (Supplemental Figure S2G); however, minimal differences are detected between canonical pathway activity, suggesting these differences may have a limited functional effect (Supplemental Figure S2H). Together, these findings reinforce previously described differences between IL-4/IL-13 and IL-10 polarized macrophages. They further suggest M2a macrophage polarization methods (IL-4/IL-13) induce slightly different gene expression patterns, and that when compared to unpolarized M0 macrophages, specific canonical pathway activity may be different between these methods.

Despite the acceptance of M2 macrophage subtypes, little work has been conducted to investigate differences between M1 macrophage populations in either *in vivo* or *in vivo* systems, based on polarization method. Researchers have induced M1 polarization through numerous methods, including exposure to LPS, IFN-γ, or a combination of LPS+IFN-γ^4,5,49,60^. However, these macrophages have largely been considered equivalent. These polarization methods have been used interchangeably and without specific thought due to the similar expression of key cell surface markers including CD80 and CD86, expression of key genes including *NOS2* and *PTGS2*, and secretion of inflammatory cytokines including IL6, IL8, and TNF with minimal further characterization^4,5,11,61^. Interestingly, each polarization method induced significantly increased expression of *CD80*, *CD86*, *NOS2, PTGS2,* and *TNF* compared to M0 macrophages; however, IFN-γ polarized macrophages consistently demonstrated the lowest fold increase in expression (data not shown). Surprisingly, IFN-γ exposed cells did not exhibit significantly increased expression of *IL6* or *IL8*, while expression of both genes varied between polarization methods with only co-exposed cells demonstrating log2 fold change expression greater than 2 (data not shown). As previous studies have demonstrated disconnects between cytokine gene expression levels and subsequently measured cytokine protein levels^62,63^, *IL6, IL8,* and *TNF* gene expression levels are likely poor markers of macrophage polarization.

We found that, after characterizing total gene expression activity, these forms of M1 macrophages demonstrate vastly different gene expression patterns, and despite similar directionality, distinct magnitudes of canonical pathway activity (Figure 3). A similar finding was suggested in a mouse model of M1 macrophages when bone marrow derived macrophages polarized with LPS+IFN-γ demonstrated similar pathway activity with differing magnitudes compared to LPS polarized peritoneal macrophages^64^. When comparing polarization methods to each other, further differences were observed. Exposures to LPS or IFN-γ induce a large degree of unique DEGs (Supplemental Figure S3), further reinforcing the notion that LPS and IFN-γ are not interchangeable methods for M1 polarization. Importantly, co-exposure to LPS+IFN-γ also induces unique DEGs not seen in either single exposure, demonstrating that co-exposure does not simply induce a combination of either exposure. When considering canonical pathway activity, LPS+IFN-γ most resembles a midpoint between LPS and IFN-γ. Pathway z-scores are generally higher than IFN-γ alone but lower than LPS alone (Supplemental Figure S3F). LPS induced the greatest degree of DEGs associated with pro-inflammatory cytokine secretion such as IL17, IL-10, and pathogen induced cytokine storm pathways (Supplemental Figure S3F). In contrast, IFN-γ induced much lower activity of these cytokine pathways while co-exposure induced a midpoint between each single exposure (Supplemental Figure S3F). Importantly, LPS+IFN-γ induced expression of the macrophage classical activation pathway to a similar degree to LPS while demonstrating much greater activity compared to IFN-γ, suggesting LPS and LPS+IFN-γ induces hallmark M1 macrophage-associated activity while IFN-γ is insufficient to induce activity commonly associated with M1 macrophages (Supplemental Figure S3F). These findings reinforce the need for researchers to consider the goals of their study and to determine the optimal polarization method to ensure their selection does not inappropriately prime or inhibit experimental models for specific responses.

While these approaches uncovered unique DEGs and differences in canonical pathway activity based on different polarization methods, it remained unclear whether these transcriptomic differences were enough to fully differentiate M1 macrophage function between polarization methods. Using the random forest machine learning algorithm^33,34^, we were able to classify macrophages based on polarization method with a high degree of accuracy, sensitivity, specificity, and precision regardless of the number of genes used for model training (Supplemental Figure S4, Figure 4). While out of bag (OOB) error rates demonstrated improvement following restriction of input genes, the initial OOB error rate of 1.03% suggested little room for improvement was available or necessary. However, as the model can be reduced to 1000 genes without demonstrating a loss in model performance but instead demonstrating slight improvements, a comparatively limited number of genes are highly impactful with clear and pronounced differences between populations. This is further demonstrated when considering the expression pattern of these genes as well as the resulting principal component analysis (PCA) clustering (Figure 5). As with the random forest modeling, macrophages sort well based on polarization method using either partition around medoid (PAM) (Figure 5A) or PCA clustering (Figure 5B). While overall patterns are more similar to each other than to unpolarized M0 macrophages, distinct patterns are clearly visible beyond initial overlaps.

In order to test the pathways impacted by these top 1000 genes and to directly compare the impact on these pathways between polarization methods, we employed gene set variation analysis (GSVA)^44^ of significantly enriched Gene Ontology (GO) biological process (BP) pathways^65^ to test pathway enrichment in each population. After collapsing gene sets into single scores, the majority of upper-level pathway enrichment was different between methods as demonstrated by significantly different GSVA scores (Figure 6). In addition, LPS+IFN-γ exposed macrophages demonstrate significantly different pathway enrichment compared to either LPS or IFN-γ exposure alone while still demonstrating similarities to each single exposure (Figure 6). These finding, along with those demonstrated by differential gene expression (Figure 3) and clustering analysis (Figure 5), fully describe M1 macrophage polarization as a complex, stimuli specific response.

We sought a more directly translatable assessment of *in vitro* M1 macrophage polarization subtypes through our previously generated random forest models (Supplemental Figure S4, Figure 4). As LPS, IFN-γ, and LPS+IFN-γ exposure induced clearly differentiable populations through random forest modeling, we sought to classify gene expression patterns of macrophages exposed to various bacterial and viral agents as LPS-, IFN-γ-, or LPS+IFN-γ-like. Most importantly, we found that virally exposed macrophages resembled unpolarized M0 or LPS+IFN-γ exposed macrophages, but not LPS or IFN-γ exposed macrophages (Figure 7). This was true regardless of the number of genes used for prediction, fully supporting the conclusion that neither LPS nor IFN-γ are appropriate models for simulating macrophage activity following exposure to viral pathogens. Indeed, the prevalence of M0 macrophage predictions further suggests that viral exposures are poorly captured using *in vitro* macrophage models, or that previously applied viral infection models have been ineffective in the context of inducing significant stimulation in *in vitro* macrophage models. In contrast, our random forest modeling suggests LPS+IFN-γ is the most widely applicable model for simulating bacterial exposures. While LPS alone was reflective of many *Listeria* and *M. tuberculosis* exposures, especially as the model was refined to include increasingly impactful genes by permutation importance (Figure 7B, Figure 7C), LPS+IFN-γ was also frequently predicted, suggesting LPS+IFN-γ may be the best model for pro-inflammatory macrophage activation following pathogen exposure. Together, these data support the notion that *in vitro* modeling of macrophages infected with pathogens needs to carefully consider the pathogen to be studied.

There are several complications with our dataset which affect our analysis and may impact our conclusions. While the use of samples from hundreds of experimental series reduces the likelihood of an individual researcher introducing significant error, it does introduce batch effects which must be considered^66^. Here, we applied the limma function removeBatchEffect which subtracts the batch effect from the data^27^, or included the batch in the regression analysis as a covariate for differential expression analysis. In addition to these general batch effects, we did not account for differences in polarization method beyond the specific stimuli employed. Specifically, differences in the concentration of stimuli, the length of stimulus, or the time between stimuli introduction and downstream analysis were not considered. However, use of dendrogram clustering is expected to remove samples which demonstrate major differences in gene expression patterns which should capture samples which received stimuli significantly outside of the norm. Finally, we did not account for the use of M-CSF versus GM-CSF for monocyte to macrophage differentiation. M-CSF has previously been demonstrated to induce an M2-like macrophage population^49,67^ while GM-CSF has been associated with M1-like states^67^. However, previous research has demonstrated the addition of conventional polarization stimuli such as LPS+IFN-γ polarizes M-CSF differentiated macrophages to M1 states^68^, suggesting the final polarization stimuli is the driving force for polarization and has a greater impact on gene expression patterns than the differentiation stimuli.

Together, our findings reinforce previous conclusions regarding macrophage polarization between unpolarized M0, pro-inflammatory M1, and pro-resolutory M2 states. Our findings further describe differences between the M2a and M2c subpopulations and suggest polarization stimuli used to generate M2a macrophages may not be entirely interchangeable. Finally, we demonstrate that, similar to M2 macrophages, M1 macrophages demonstrate clear differences in gene expression patterns following stimuli with commonly used polarization regimens. These findings were generated from a significantly larger dataset than previously employed for similar studies and compiles RNAseq data generated from hundreds of individual study series, offering a wide-ranging dataset which comprehensively represents macrophage populations without the concerns generated from single-source studies. While additional studies are required to directly compare the functional activity of pro-inflammatory macrophages following polarization with specific stimuli, these findings identify the need for researchers to consider polarization method when designing and analyzing *in vitro* macrophage studies.

## Acknowledgements

The authors would like to acknowledge the University of North Carolina at Chapel Hill Curriculum in Toxicology and Environmental Medicine Leon and Bertha Golberg Postdoctoral Fellowship for training support.

## Data Availability

All RNAseq count data used in this study were obtained from ARCHS^4^ version 2.2, accessed on October 13^th^, 2023 at ^19^. R code used for data analysis and figure generation is available at: https://github.com/UNC-CEMALB/Comprehensive-Unbiased-RNAseq-Database-Uncovers-New-Human-Monocyte-Derived-Macrophage-Subtypes-.

## Contributions

TS and IJ conceived and designed the study. TS acquired, analyzed, and interpreted all data. EH and JR assisted in review and refinement of the study design and manuscript. AP assisted in final review of R code. TS and IJ drafted the manuscript. IJ was project leader. All authors read and approved the final manuscript.

## Funding

This research was funded by the National Institute of Environmental Health Sciences (NIEHS) T32 ES007126 and R01 ES031173.

## Disclosures

The authors have no conflicts of interest to disclose.

## Notes

### Competing Interest Statement

The authors have declared no competing interest.

