## Supplemental Materials for "Leveraging a Comprehensive Unbiased RNAseq Database Uncovers New Human Monocyte-Derived Macrophage Subtypes Within Commonly Employed *In Vitro* Polarization Methods"

### Supplemental Results

*M2 Macrophages Demonstrate Slight Polarization Method-Dependent Gene Expression Patterns*

When differential expression is assessed between polarization methods, negligible differences are detected between IL-4 and IL-13 populations (Supplemental Figure S2A) with a total of 18 DEGs. In contrast, IL-10 polarization induced clear difference in expression patterns compared to IL-4 exposed cells with 283 DEGs (Supplemental Figure S2B), but not IL-13 exposed cells with only 33 DEGs (Supplemental Figure S2C). Compared to these single polarization stimuli, co-exposure to IL-4+IL-13 induced 76 DEGs compared to IL-4 exposure alone (Supplemental Figure S2D) and 17 DEGs compared to IL-13 alone (Supplemental Figure S2E), suggesting IL-13 slightly potentiates the effects of IL-4 polarization, but that IL-4 has a minimal impact on modifying IL-13 induced polarization. In contrast, major differences were detected between IL-4+IL-13 co-exposed macrophages and IL-10 exposed macrophages, with 240 DEGs detected (Supplemental Figure S2F). While a large degree of overlap was detected between DEGs in IL-4 and IL-4+IL-13 co-exposed macrophages compared to polarization with IL-10 (Supplemental Figure S2G), a similar degree of DEGs were unique to each comparison. Together these data suggest that IL-10 induces a clearly different macrophage state compared to IL-4 and IL-4+IL-13, but that IL-4 and IL-4+IL-13 may also demonstrate distinct gene expression patterns to each other. While IPA demonstrated limited activation or inhibition of canonical pathways overall, the majority of pathway modifications were detected between cells exposed to IL-10 versus IL-4, IL-13, or a combination of IL-4+IL-13 (Supplemental Figure S2H). These findings further support previous findings suggesting that IL-4+IL-13 induces polarization of a distinct subset of M2 macrophages compared to IL-10 polarized M2 macrophages but leaves open the possibility that IL-4+IL-13 co-exposure may induce slight differences in macrophage populations compared to IL-4 exposure alone.

*M1 Macrophage Polarization Method Induces Distinct Gene Expression Patterns Compared to Other Polarization Methods*

As with M2 macrophages, when comparing polarization methods to each other, single exposure to IFN-γ or LPS induces 503 DEGs between subsets (Supplemental Figure S3A, S3B) while co-exposure to LPS+IFN-γ induces 1,094 DEGs compared to IFN-γ exposure (Supplemental Figure S3C) and 730 DEGs compared to LPS exposure alone (Supplemental Figure S3D). Interestingly, a major overlap of 385 DEGs was detected between co-exposed macrophages compared to either single exposure (Supplemental Figure S3E), suggesting co-exposure to LPS+IFN-γ activates unique genes not induced by either polarization stimuli alone. However, co-exposure does not induce expression of all genes induced by single exposures, as 391 DEGs are unique to the IFN exposed group and 368 DEGs in the LPS exposed group (Supplemental Figure S3E). These findings can be further examined through inspection of canonical pathway activity (Supplemental Figure S3F). LPS exposed groups demonstrated clear induction of numerous pro-inflammatory pathways traditionally associated with M1 macrophages including pro-inflammatory cytokine secretion and classical macrophage activation (Supplemental Figure S3F). Of the top pathways, co-exposure to LPS+IFN-γ largely blunted pathway modifications compared to LPS exposure alone but demonstrated greater macrophage classical activation signaling and lower macrophage alternative activation signaling, suggesting co-exposure induces gene expression patterns closest to the literature definition of M1 macrophage (Supplemental Figure S3F). Interestingly, of the top pathways, only LXR/RXR activation is elevated in IFN-γ compared to LPS exposed cells, however, co-exposure to LPS+IFN-γ demonstrated even greater activation compared to IFN-γ exposure alone (Supplemental Figure S3F). These findings together suggest LPS and IFN-γ are capable of both blunting and potentiating the effects of the other, indicating that co-exposed macrophages are a wholly unique macrophage population rather than a combination of each single exposure.

### Supplemental Figure Legends

**Supplemental Figure S1. Human Monocyte-Derived Macrophages Demonstrate High Within Polarization Method Correlations Following Sample Selection and Batch Selection.** Density plots demonstrating human monocyte-derived macrophage sample correlations within indicated groups composed of single polarization methods. Red lines represent all sample before sample selection. Blue lines represent sample correlations following dendrogram clustering and removal of samples. Green lines represent sample correlations after sample selection and batch correction. Black vertical lines represent the sample correlation cutoff of 0.75.

**Supplemental Figure S2. M2 Macrophages Demonstrate Slight Polarization Method-Specific Gene Expression Patterns.** A-F. Volcano plots representing results of differential expression analyses between indicated groups. G. Euler plots describing overlap of significantly differentially expressed genes between groups. H. Ingenuity Pathway Analysis (IPA) canonical pathway activation z-score heatmap. Significance cutoffs of Benjamini-Hochberg (BH)-adjusted P-values $< 0.05$ and absolute $\log_2$ fold changes $\geq 2$ were employed for all analyses. NA values represent pathway z-scores which could not be calculated.

**Supplemental Figure S3. M1 Macrophages Demonstrate Major Polarization Method-Specific Gene Expression Patterns.** A-D. Volcano plots representing results of differential expression analyses between indicated groups. E. Euler plots describing overlap of significantly differentially expressed genes between groups. F. Ingenuity Pathway Analysis (IPA) canonical pathway activation z-score heatmap. Significance cutoffs of Benjamini-Hochberg (BH)-adjusted P-values $< 0.05$ and absolute $\log_2$ fold changes $\geq 2$ were employed for all analyses. NA values represent pathway z-scores which could not be calculated.

**Supplemental Figure S4. Restriction of Input Genes Has a Minimal Effect on Random Forest Model Performance.** A. Random Forest model generated using all genes (34,479 genes) demonstrating adjusted P-values < 0.05 following one-way ANOVA with Benjamini-Hochberg (BH) procedure conducted between M0, LPS, IFN-γ, and LPS+IFN-γ groups. B. Random Forest model generated using the top 10,000 genes from the unrestricted model as determined by permutation importance. Random Forest confusion matrices (top) describing the number of samples from the test data subset predicted as each class versus actual class. Model performance metrics (middle, bottom) describe the ability of the model to correctly classify samples and to avoid misclassifying samples across classes.

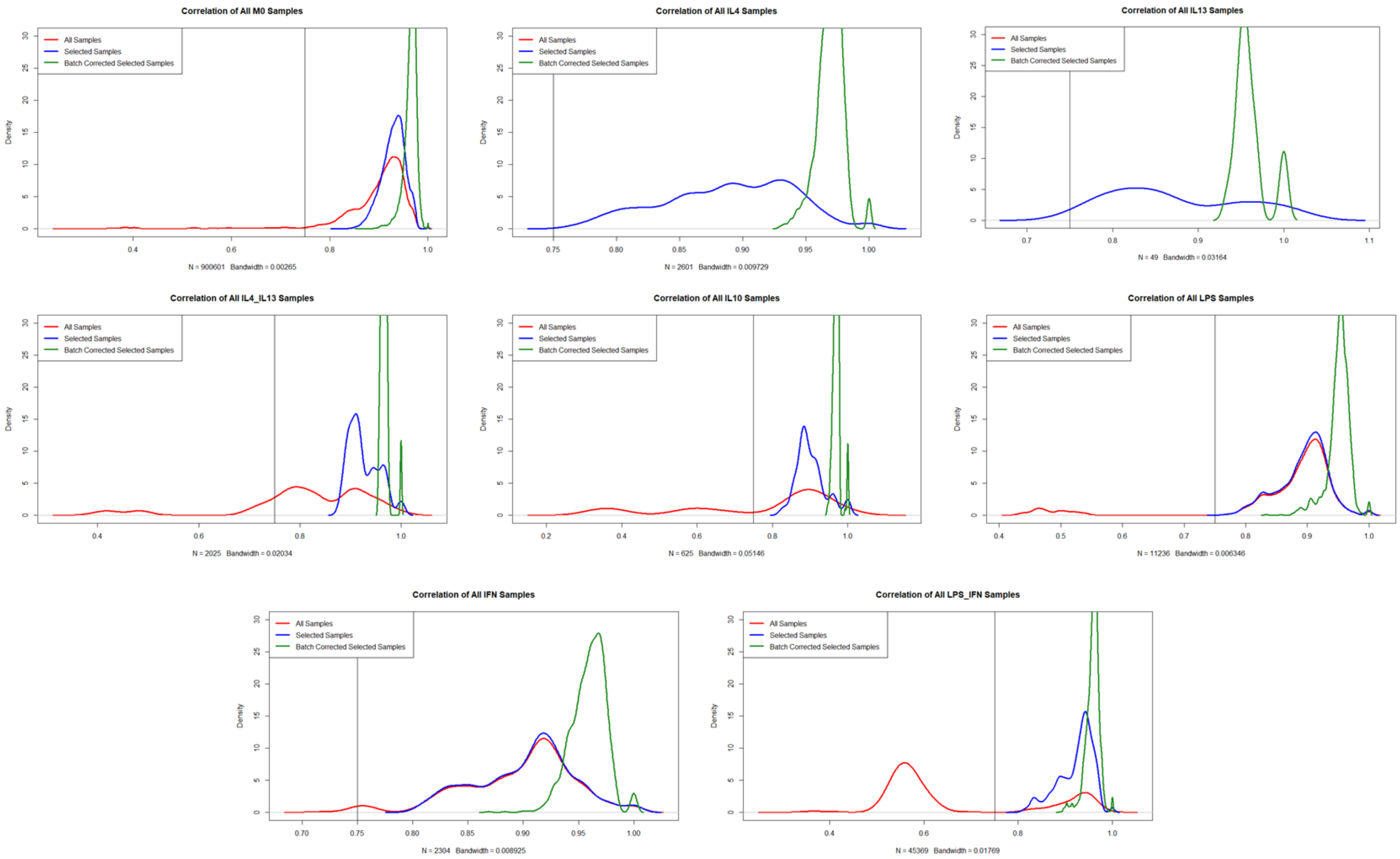


**Supplemental Figure S1.**

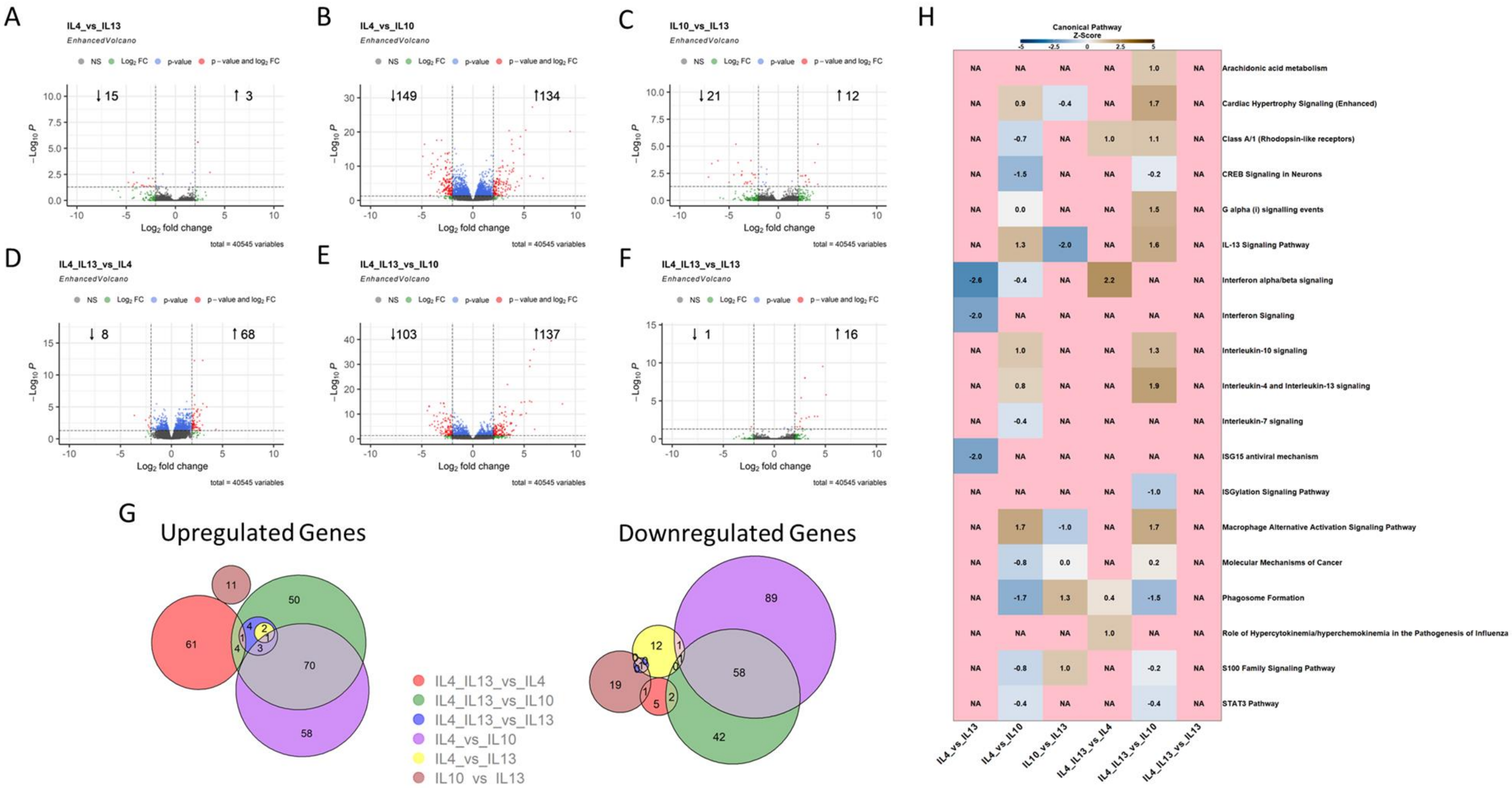


**Supplemental Figure S2.**

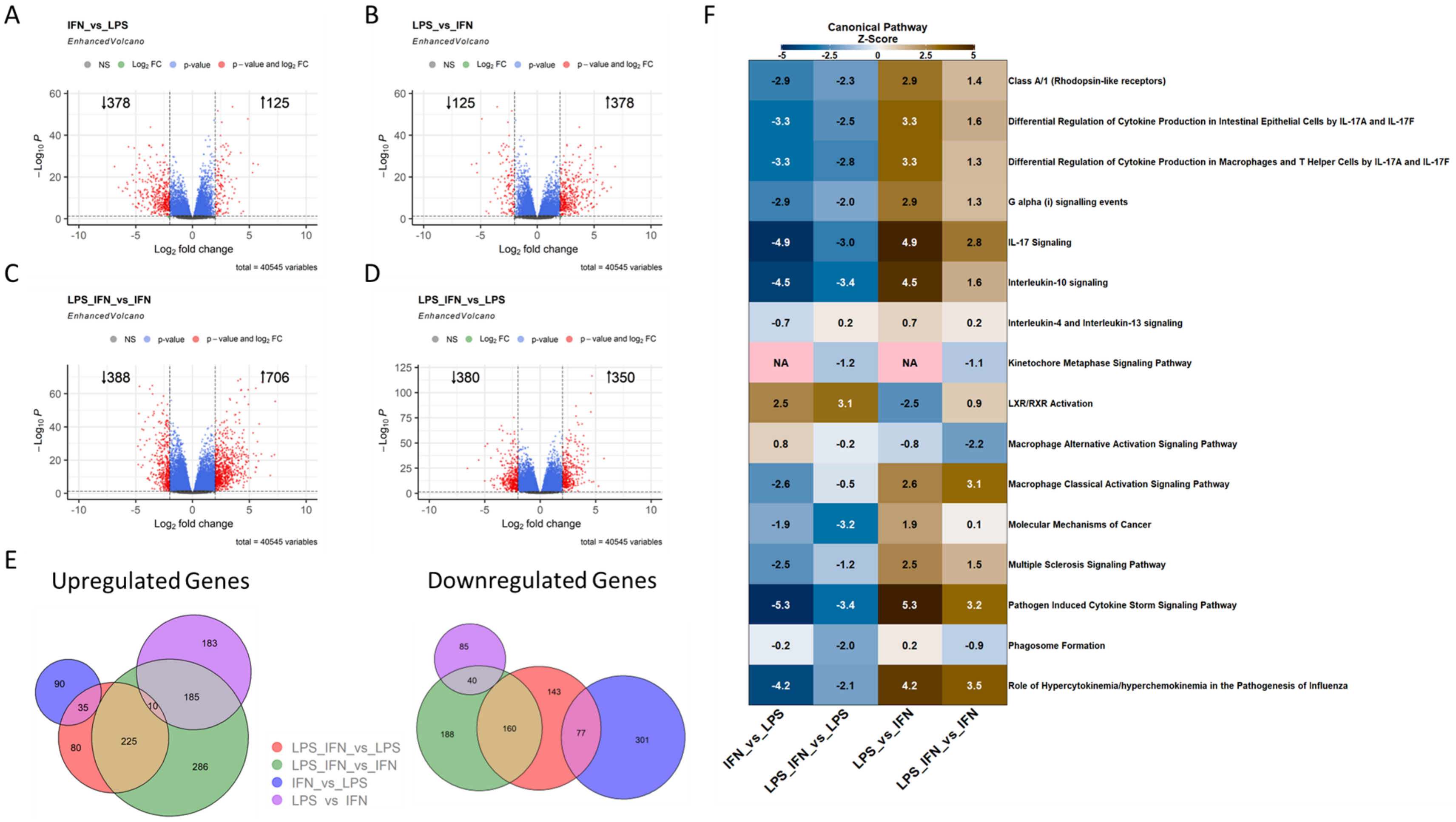


**Supplemental Figure S3.**

A

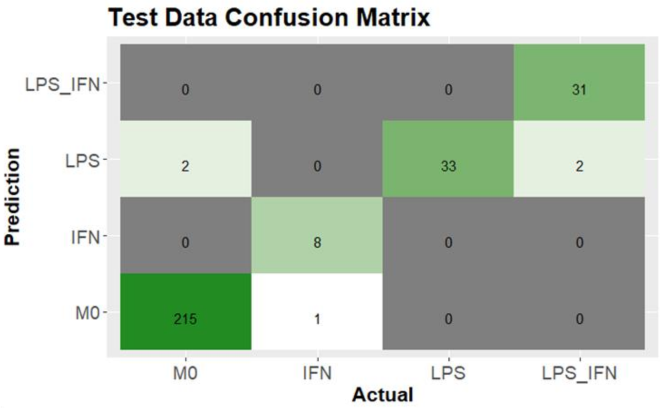


| Accuracy | Kappa | AccuracyLower | AccuracyUpper | AccuracyNull |
|---|---|---|---|---|
| 0.9829 | 0.9595 | 0.9605 | 0.9944 | 0.7432 |

| | Sensitivity | Specificity | Pos.Pred.Value | Neg.Pred.Value | Precision | Recall | F1 |
|---|---|---|---|---|---|---|---|
| *Class: M0* | 0.99 | 0.99 | 1 | 0.97 | 1 | 0.99 | 0.99 |
| *Class: IFN* | 0.89 | 1 | 1 | 1 | 1 | 0.89 | 0.94 |
| *Class: LPS* | 1 | 0.98 | 0.89 | 1 | 0.89 | 1 | 0.94 |
| *Class: LPS_IFN* | 0.94 | 1 | 1 | 0.99 | 1 | 0.94 | 0.97 |

B

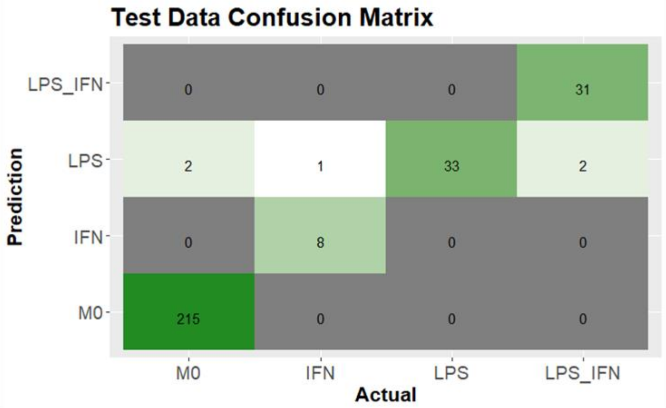


| Accuracy | Kappa | AccuracyLower | AccuracyUpper | AccuracyNull |
|---|---|---|---|---|
| 0.9829 | 0.9597 | 0.9605 | 0.9944 | 0.7432 |

| | Sensitivity | Specificity | Pos.Pred.Value | Neg.Pred.Value | Precision | Recall | F1 |
|---|---|---|---|---|---|---|---|
| *Class: M0* | 0.99 | 1 | 1 | 0.97 | 1 | 0.99 | 1 |
| *Class: IFN* | 0.89 | 1 | 1 | 1 | 1 | 0.89 | 0.94 |
| *Class: LPS* | 1 | 0.98 | 0.87 | 1 | 0.87 | 1 | 0.93 |
| *Class: LPS_IFN* | 0.94 | 1 | 1 | 0.99 | 1 | 0.94 | 0.97 |

**Supplemental Figure S4.**
